# Pre- and postnatal maternal depressive symptoms associate with local connectivity of the left amygdala in 5-year-olds

**DOI:** 10.1101/2024.06.21.600007

**Authors:** Elena Vartiainen, Anni Copeland, Elmo P. Pulli, Venla Kumpulainen, Eero Silver, Olli Rajasilta, Ashmeet Jolly, Silja Luotonen, Hilyatushalihah K. Audah, Niloofar Hashempour, Wajiha Bano, Ilkka Suuronen, Ekaterina Saukko, Suvi Häkkinen, Hasse Karlsson, Linnea Karlsson, Jetro J. Tuulari

**Affiliations:** FinnBrain Birth Cohort Study, Turku Brain and Mind Center, Department of Clinical Medicine, University of Turku, Turku, Finland; Centre for Population Health Research, Turku University Hospital and University of Turku; Neurocenter, Turku University Hospital, Turku, Finland; Department of Psychology and Speech Language Pathology, University of Turku, Turku, Finland; Department of Teacher Education, University of Turku, Turku, Finland; Department of Psychiatry, Turku University Hospital and University of Turku, Turku, Finland; Department of Radiology, Turku University Hospital and University of Turku, Turku, Finland; Department of Neuroscience, University of California, Berkeley, Berkeley, CA, USA; Department of Child Psychiatry, Turku University Hospital, Turku, Finland; Department of Public Health, University of Turku and Turku University Hospital; Clinical Neurosciences, University of Turku, Turku, Finland

**Keywords:** brain development, fMRI, maternal depression, amygdala, ReHo, children

## Abstract

**Background:** Maternal depressive symptoms can influence brain development in offspring, prenatally through intrauterine programming, and postnatally through caregiving related mother–child interaction.

**Methods:** The participants were 5-year-old mother-child dyads from the FinnBrain Birth Cohort Study (N = 68; 28 boys, 40 girls). Maternal depressive symptoms were assessed with the Edinburgh Postnatal Depression Scale (EPDS) at gestational week 24, 3 months, 6 months, and 12 months postnatal. Children’s brain imaging data were acquired with task-free functional magnetic resonance imaging (fMRI) at the age of 5 years in 7 min scans while watching the *Inscapes* movie. The derived brain metrics included whole brain regional homogeneity (ReHo) and seed-based connectivity maps of the bilateral amygdalae.

**Results:** We found that maternal depressive symptoms were positively associated with ReHo values of the left amygdala. The association was highly localised and strongest with the maternal depressive symptoms at three months postnatal. Seed-based connectivity analysis did not reveal associations between distal connectivity of the left amygdala region and maternal depressive symptoms.

**Conclusions:** These results suggest that maternal depressive symptoms soon after birth may influence offspring’s neurodevelopment in the local functional coherence in the left amygdala. They underline the potential relevance of postnatal maternal distress exposure on neurodevelopment that has received much less attention than prenatal exposures. These results offer a possible thus far understudied pathway of intergenerational effects of perinatal depression that should be further explored in future studies.

## 1. Introduction

Maternal psychological distress in pregnancy can affect fetal brain development in fundamental ways (1,2). The prevalence of clinically relevant level of depressive symptoms, including major and minor depressive disorder, in pregnancy is around 11% in the first trimester and 8.5% in the second and third trimester (3). Neuroimaging studies using structural and diffusion MRI studies have indicated that maternal prenatal stress, including depression, is associated with changes in child brain morphology (2,4,5). These findings affirm that the prenatal period is critical for fetal neurodevelopment and poor maternal well-being during pregnancy may have long-lasting effects on the offspring. Effects of prenatal depressive symptoms are found in brain structures that are responsible for e.g. emotional regulation, cognition, memory, and decision-making – with a majority of the studies implication the amygdala (6).

Resting-state functional magnetic resonance imaging (rs-fMRI) can provide novel information on maternal distress as a predictor of offspring’s neurodevelopment (7–10). An rs-fMRI study on infant neurodevelopment showed that maternal prenatal depressive symptoms were associated with increased functional connectivity between medial prefrontal cortex (mPFC) and left amygdala in 6-month-old infants (9). Rajasilta et al. (2023) demonstrated that in infants approximately one moth of age maternal prenatal distress was positively to rs-fMRI-derived fractional amplitude of low-frequency fluctuations (fALFF) in mPFC which indicates that prenatal stress may affect functional features of the maturing brain during gestation (11). Children of mothers who experienced psychological distress late in their pregnancy had increased amygdala functional connectivity with the ventromedial PFC and anterior insula (12). In comparison to prenatal maternal depressive symptoms, early postnatal maternal wellbeing and later child development have received significantly less attention in neuroimaging studies, even though the first years of life are crucial time for parent-child bonding and child development.

To the best of our knowledge, there is only one prior functional neuroimaging study linking both pre- and postnatal maternal depressive symptoms to adverse child brain connectivity outcomes in early childhood. This study by Soe et al. (2018) investigated the relationship between perinatal maternal depressive symptoms and bilateral amygdala in 4.5-year-old children. They found significant positive associations between maternal prenatal depressive symptoms and the left amygdala functional connectivity with the right insula and putamen, bilateral subgenual anterior cingulate cortex (ACC) and caudate. Similar associations were found between the right amygdala and left orbitofrontal cortex and temporal pole. Greater pre-than postnatal depressive symptoms had associations with lower functional connectivity of the left amygdala with the bilateral subgenual ACC and left caudate. Correspondingly, greater prenatal maternal depressive symptoms were associated with lower functional connectivity of the right amygdala with the left orbitofrontal cortex, insula, and caudate. These findings were gender-specific to girls. However, the study did not find significant interactions of gender with pre- or postnatal maternal depressive symptoms on amygdala functional connectivity (13).

In the current study, we investigate whether maternal pre- and postnatal depressive symptoms associate with 5-year-olds’ local connectivity across the whole brain as assessed by regional homogeneity (ReHo), a measure derived from task-free fMRI while the participants were watching *Inscapes* movie (14). Second, replicating the analyses in prior work (13), we mapped the association of maternal depressive symptoms to bilateral amygdala seed-based connectivity. Third, multiple regression analyses were performed to assess the associations between maternal depressive symptoms and bilateral amygdala ReHo values. This was an exploratory study and thus no explicit hypotheses were placed regarding the direction or strength of the associations.

## 2. Methods and Materials

This study was conducted in accordance with the Declaration of Helsinki, and it was approved by the Joint Ethics Committee of the University of Turku and the Hospital District of Southwest Finland (15.03.2011) §95, ETMK: 31/180/2011. Written informed consent was obtained from the participants, and parents gave consent on behalf of their children. We followed the Strengthening the Reporting of Observational studies in Epidemiology (STROBE) reporting guidelines for cohort studies (https://www.strobe-statement.org). The participant criteria, MRI scanning visits, image acquisition and preprocessing are identical to those used in our prior work (15).

### 2.1 Participants

The participants are part of the FinnBrain Birth Cohort Study, which prospectively examines the influence of genetic and environmental factors on child development and later mental and physical health outcomes (16). Pregnant women attending their first trimester ultrasound were recruited in maternal welfare clinics in the Turku region of the Southwest Finland Hospital District and the Åland Islands between December 2011 and April 2015. Ultrasound-verified pregnancy and a sufficient knowledge of Finnish or Swedish were required for participation.

The exclusion criteria for the neuroimaging study were: (1) born before gestational week 35 (week 32 for those with exposure to maternal prenatal synthetic glucocorticoid treatment), (2) developmental or major organ abnormalities in senses or communication (e.g., blindness, deafness, congenital heart disease), (3) known long-term medical diagnosis (e.g., epilepsy, autism), (4) ongoing medical examinations or clinical follow-up in a hospital, (5) the child using continuous, daily medication (including oral medications, topical creams, and inhalants; desmopressin was allowed), (6) history of head trauma (defined as concussion necessitating clinical follow up in a health care setting), (7) metallic ear tubes, and (8) routine MRI contraindications.

In total, 203 children participated in a neuroimaging visit at 5 years of age, and 77 of them had successful functional scans due to limited subject compliance (the fMRI data was acquired last), out of which 68 had the maternal Edinburgh Postnatal Depression Scale (EPDS) questionnaires collected, and were included in the study after quality control steps outlined below. Participant characteristics are reported in Table 1.

**Table 1.** Demographics of the study participants.

|  | **Valid** | **Missing** | **Mean** | **SD** | **Range** |
| --- | --- | --- | --- | --- | --- |
| Child age at scan (years) | 68 | 0 | 5.39 | 0.12 | 0.51 |
| Duration of gestation (weeks) | 68 | 0 | 39.84 | 1.34 | 6.43 |
| Child birthweight (grams) | 68 | 0 | 3551 | 430 | 2370 |
| Maternal age at childbirth (years) | 68 | 0 | 30.40 | 4.63 | 21.00 |
| Maternal pre-pregnancy BMI (kg/m<sup>2</sup>) | 68 | 0 | 23.52 | 3.99 | 19.25 |
| EPDS sum score at gestational week 24 | 65 | 3 | 5.12 | 4.23 | 21.00 |
| EPDS sum score at 3 months postnatal | 68 | 0 | 3.97 | 3.52 | 12.22 |
| EPDS sum score at 6 months postnatal | 60 | 8 | 5.19 | 4.63 | 19.00 |
| EPDS sum score at 12 months postnatal | 57 | 11 | 4.39 | 4.32 | 21.00 |
| SCL-90 sum score at gestational week 24 | 65 | 3 | 3.41 | 3.55 | 14.00 |
| SCL-90 sum score at 3 months postnatal | 68 | 0 | 2.00 | 3.12 | 15.00 |
| SCL-90 sum score at 6 months postnatal | 60 | 8 | 3.21 | 4.75 | 24.00 |

| **Child sex** | **Frequency** | **Percent** |
| --- | --- | --- |
| Boy | 28 | 41.2% |
| Girl | 40 | 58.8% |
| Missing | 0 | 0 |

| **Maternal education** | **Frequency** | **Percent** |
| --- | --- | --- |
| Upper secondary school or vocational school or lower | 14 | 20.6% |
| University of applied sciences | 17 | 25.0% |
| University | 36 | 52.9% |
| Missing | 1 | 1.5% |

| **Maternal cultural background** | **Frequency** | **Percent** |
| --- | --- | --- |
| Finnish | 66 | 97.1% |
| Other | 2 | 2.9% |
| Missing | 0 | 0 |

| **Smoking during pregnancy** | **Frequency** | **Percent** |
| --- | --- | --- |
| No smoking | 64 | 94.1% |
| During first trimester | 2 | 2.9% |
| During third trimester | 2 | 2.9% |
| Missing | 0 | 0 |

SD, standard deviation; BMI, body mass index; EPDS, Edinburgh Postnatal Depression Scale; SCL-90, Symptom Checklist 90

### 2.2 Maternal depressive and anxiety symptoms and perinatal data

Questionnaires assessing maternal psychological health were filled in by the mothers in gestational week 24 and 3 months, 6 months, and 12 months postnatal. We assessed prenatal depressive symptoms at gestational week 24 because at this timepoint neurogenesis takes place, cortex begins to fold and myelin starts to develop (17–19). Maternal depressive symptoms were assessed with the EPDS (20), which has been validated for use during pregnancy. This 10-item questionnaire is scaled from 0 to 30 points with a higher score denoting increased symptom severity. A score of 10 has been implicated as a clinically meaningful threshold for symptoms of depression in pregnancy (21). Pre- and postnatal maternal anxiety symptoms were evaluated using the anxiety scale of Symptom Checklist 90 (SCL-90) (22) at gestational week 24, 3 months and 6 months postnatal. The anxiety subscale of SCL-90 is a reliable and valid measure of anxiety symptoms in both clinical and research settings and the questionnaire consists of 10 items scaled from 0 to 4. Obstetric data was retrieved from the Finnish National Birth Register (National Institute for Health and Welfare, www.thl.fi). Other information was gathered using questionnaires on behalf FinnBrain Birth Cohort Study.

### 2.3 Magnetic resonance imaging visits

All MRI scans were performed for research purposes, and participants were scanned awake and without sedation. The imaging was performed at the Department of Radiology, Turku University Hospital between October 2017 and March 2021. The practical arrangements to perform child friendly visits, more details of the visit have been described previously (15). Anatomical images were screened by an experienced neuroradiologist for incidental findings. None of the participants included in the present study had clinically relevant incidental findings.

### 2.4 Image acquisition

The MRI scans were conducted on a Siemens Magnetom Skyra fit 3T scanner (Siemens Medical Solutions, Erlangen, Germany). A 20-element Head/Neck Matrix coil allowed the use of the generalized autocalibrating partially parallel acquisition (GRAPPA) technique to accelerate acquisitions (parallel acquisition technique [PAT] factor of 2). The scans included a high-resolution T1 magnetization-prepared rapid gradient echo (MPRAGE), a T2 turbo-spin echo (TSE), diffusion tensor imaging (DTI), and a 7-min fMRI. The fMRI consisted of 170 volumes with voxel size 3.0 × 3.0 × 3.0 mm^3^, TR 2500 ms, TE 30.0 ms, flip angle of 80°, and 42 axial slices without gaps. Full cerebellar coverage was not possible in all participants. Prior to fMRI acquisition, all children had rested by watching a movie or a TV show of their choice during the 30–45 min required for structural scanning.

If the child had fallen asleep, they were gently awakened. During the fMRI sequence, participants were instructed to stay as still as possible with their eyes open. To minimize motion and reduce cognitive load, the *Inscapes* movie was played during fMRI data collection (14). Visual stimuli were presented on an MRI-compatible 32′′ LCD monitor with full HD resolution (Nordic Neuro Lab) located at the foot of the bed of the scanner, which participants could watch via mirrors mounted on the head coil. The total scanning time was limited to 1 hour, and the imaging was discontinued if the child expressed unwillingness to continue at any point.

### 2.5 Image preprocessing and estimating regional homogeneity (ReHo)

Functional MRI data were slice-timing corrected and motion corrected in the FMRIB Software Library (v6.00, FSL; Jenkinson et al., 2012) v6.00 relative to a manually chosen reference volume, free of major artifacts. Motion outliers were estimated using artifact detection tools (ART) (https://www.nitrc.org/projects/artifact_detect). We tagged images as outliers if they had composite motion threshold > 2 mm or DVARS > 9, which are default parameters in the ART toolbox. All children included in the final statistical analyses had a full fMRI sequence of 170 volumes, and a maximum of 50 volumes were tagged as outliers by ART. The descriptive statistics for motion were as follows (of full sample N = 68): motion outliers (mean 15, range 0–49), mean absolute displacement (mean 0.73, range 0.06–3.71, mm), and mean relative displacement (mean 0.25, range 0.02–1.36, mm). Anatomical masks for white matter and cerebrospinal fluid were defined in the Montreal Neurological Institute (MNI) standard space and spatially normalized using FSL, then registered to functional data with an affine transformation. Average signal in white matter and cerebrospinal fluid as well as 24 motion covariates (the six realignment parameters and their temporal derivatives and quadratic terms) were included as nuisance covariates. Taken together, denoising consisted of outlier rejection, nuisance regression, detrending, and high-pass filtering (0.008 Hz).

Our main imaging derivative was regional homogeneity (ReHo), a data-driven measure of local voxel-wise connectivity. ReHo describes synchronization between timeseries of a given voxel in its neighbours based on Kendall’s coefficient of concordance and is interpreted as a sign of synchronised activation or deactivation in blood-oxygen-level-dependent timeseries (24). ReHo was computed as implemented in DPABI (number of voxels in a cluster; N = 27). For group analysis, ReHo maps were normalized non-linearly to 1.0 × 1.0 × 1.0 mm^3^ MNI space using FSL FNIRT. Finally, the data were smoothed with a Gaussian filter of 6 mm full width at half maximum (FWHM).

### 2.6 Seed-based connectivity analysis (SCA)

The primary analyses focused on whole brain ReHo maps. Our second goals was to replicate prior work (13) by studying the association of maternal depressive symptoms to bilateral amygdala seed-based connectivity. Additionally, in line with our prior work (15,25), we had predefined plans to conduct complementary SCAs guided by the ReHo results. The SCA analyses were performed with FSL tools using the same pre-processing and nuisance regression as for the ReHo analyses except that the interquartile range (obtained via fsl_motion_outliers) of DVARS was used for removal of motion corrupted volumes after confirming that it matched the ReHo pipeline described above.

### 2.7 Statistical analysis

Whole brain voxelwise statistical analyses for the ReHo maps and the SCA connectivity maps were performed with SPM12 software (https://www.fil.ion.ucl.ac.uk/spm/software/spm12/) with general linear models (GLM) and “multiple regression” design for ReHo. Association between maternal perinatal depressive symptoms and ReHo of the bilateral amygdala was assessed in two stages. First, child age at scan and sex were set as the independent variables (IV) of no interest and EPDS score was set as the main explanatory variable (EV). Second, we added IVs of no interest including maternal socioeconomic status (SES) measured by maternal educational level, maternal pre-pregnancy BMI, and child’s ponderal index (measured at MRI scan, relationship between body mass and height; mass/height^3^, (26)), and with maternal anxiety symptom score (SCL-90 score). GLM models were performed for EPDS scores separately for each of the time points (gestational week 24, 3 months, 6 months, and 12 months postnatal). Prior to statistical testing we used an inclusive binary grey matter mask. The *a priori* statistical threshold for clusters was set at *p* < 0.001 and secondarily at *p* < 0.005 and corrected with family-wise error (FWE) at cluster level at *p* < 0.05.

Complementary region of interest based linear regression models were performed for bilateral amygdala mean ReHo values with RStudio (R Core Team (2024). _R: A Language and Environment for Statistical Computing_. R Foundation for Statistical Computing, Vienna, Austria. https://www.R-project.org/.) This analysis was carried out to describe the effect sizes of the associations between the ReHo of the amygdala, and the independent variables in line with the SPM12 models, and with other additional independent variables described below. The ReHo values were obtained by creating binary masks of bilateral amygdala from the AAL atlas, using them to mask normalised ReHo maps and estimating mean with fslmaths. All regression models were performed for left and right amygdala mean ReHo values separately. In the first regression model, IVs were set as EPDS score (gestational week 24, 3 months, 6 months, and 12 months postnatal separately), child sex, and child age at scan. In the second model, IVs were set as EPDS score (gestational week 24, 3 months, 6 months, and 12 months postnatal separately), child sex, child age at scan, maternal SES, maternal pre-pregnancy BMI, and ponderal index. Lastly, we included maternal anxiety symptoms (SCL-90 scores at gestational week 24, 3 months, and 6 months) in the second model to test if the association was specific to depressive symptoms. Missing EPDS score, SCL-90 score, and maternal educational level values were mean imputed for statistical analyses. We checked that the regression models met basic assumptions: multicollinearity through variation inflation factor (all variance inflation factors, VIF’s < 2.3) and that the residuals were normally distributed through visual inspection of Q–Q plots and histograms of the residuals. We report uncorrected *p*-values throughout the manuscript but note that we performed eight statistical tests, 2 (left and right amygdala) x 4 (EPDS questionnaire timepoints), and the Bonferroni corrected *p*-values are *p* < 0.00625.

## 3. Results

The sample characteristics are reported in Table 1. The sample included 28 boys and 40 girls. Maternal EPDS scores showed variability across different measurement points so that the mean scores were 5.12 (gestational week 24), 3.97 (3 months postnatal), 5.19 (6 months postnatal), and 4.39 (12 months postnatal). The corresponding SCL-90 scores were 3.41 (gestational week 24), 2.00 (3 months postnatal), and 3.21 (6 months postnatal).

### 3.1 Whole brain voxelwise associations

We did not reveal positive or negative associations between EDPS sum score at gestational week 24 and child brain ReHo map of left amygdala when child sex and age at scan were controlled.

Significant positive association between mothers’ EPDS score at 3 months postnatal and child brain ReHo map of left amygdala region was found when child sex and age at scan were controlled (thresholded at *p* < 0.001, *p* = 0.038 FWE corrected, cluster size (kE) 371) (Supplemental Figure 1). The association remained significant when maternal pre-pregnancy BMI, maternal SES and ponderal index were included as covariates (thresholded at *p* < 0.001, *p* = 0.059 FWE corrected, cluster size (kE) 326) (Figure 1) but the association did not remain significant when SCL-90 score was included as a covariate in addition. This might be because of correlations between EPDS and SCL scores and indeed both depressive symptoms and depressive symptoms associated with the left amygdala ReHo values (Supplemental Table A). There were no negative associations between mothers’ EPDS score at 3 months postnatal and ReHo map of the left amygdala region.

**Figure 1.**
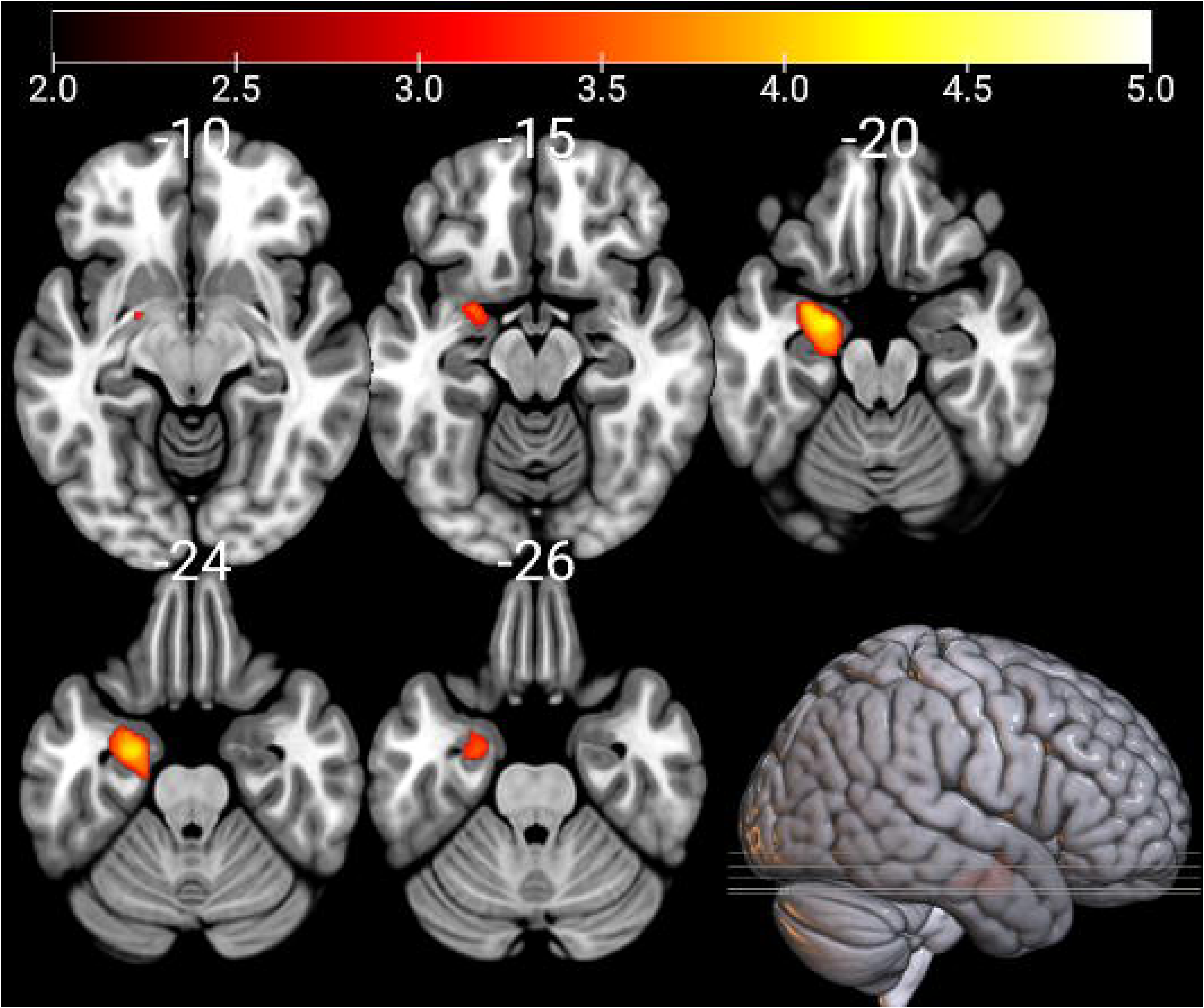
ReHo values of the left amygdala associate positively with EPDS scores at 3 months postnatal; controlling for child sex, age at scan, maternal BMI, maternal socioeconomic status, and ponderal index. The results have been thresholded at p < 0.001, FDR multiple comparisons corrected at the cluster level. The color bars depict t-values.

No statistically significant associations were found between EPDS score at 6 months postnatal and child brain ReHo map of left amygdala region with the whole brain analyses. However, we observed a positive association between mothers’ EPDS score at 12 months postnatal and ReHo map of the left amygdala region when child sex and age at scan were set as covariates (thresholded at *p* < 0.005, *p* = 0.011 FWE corrected, cluster size (kE) 1152) and when maternal pre-pregnancy BMI, maternal SES and ponderal index were controlled in addition (thresholded at *p* < 0.005, *p* = 0.028 FWE corrected, cluster size (kE) 958).

There were no negative associations between child amygdala ReHo values and maternal EPDS scores in any of the time points. We did not find significant associations in seed-based connectivity analysis of our seed region of interest, bilateral amygdala, to the rest of the brain, and the EPDS scores.

### 3.2 Region of interest analyses of ReHo for bilateral amygdala

Summary of all statistically significant results of the linear regression analyses for bilateral amygdala mean ReHo values are shown in Table 2. Additionally, summary of all linear regression analysis for left amygdala mean ReHo values are shown in Supplemental Table A and for right amygdala mean ReHo values in Supplemental Table B.

**Table 2.** Summary of the statistically significant results of the regression analyses for bilateral amygdala mean ReHo values.

| Relation | Controlled covariates | $\beta$ | $p$ |
| --- | --- | --- | --- |
| Left Amygdala ReHo and EPDS sum score at gestantional week 24 | Child sex, and age at scan | 0.25 | < 0.05 |
| Left Amygdala ReHo and EPDS sum score at 3 months | Child sex, and age at scan | 0.48 | < 0.001 |
| Left Amygdala ReHo and EPDS sum score at 3 months | Child sex, age at scan, maternal SES, maternal pre-pregnancy BMI, and ponderal index | 0.47 | < 0.001 |
| Left Amygdala ReHo and EPDS sum score at 3 months | Child sex, age at scan, maternal SES, maternal pre-pregnancy BMI, ponderal index, and SCL-90 sum score at 3 months | 0.34 | < 0.05 |
| Right Amygdala ReHo and EPDS sum score at 3 months | Child sex, and age at scan | 0.28 | < 0.05 |
| Right Amygdala ReHo and EPDS sum score at 3 months | Child sex, age at scan, maternal SES, maternal pre-pregnancy BMI, and ponderal index | 0.29 | < 0.05 |
| Left Amygdala ReHo and EPDS sum score at 6 months | Child sex, and age at scan | 0.38 | < 0.01 |
| Left Amygdala ReHo and EPDS sum score at 6 months | Child sex, age at scan, maternal SES, maternal pre-pregnancy BMI, and ponderal index | 0.37 | < 0.01 |
| Left Amygdala ReHo and EPDS sum score at 12 months | Child sex, and age at scan | 0.42 | < 0.001 |
| Left Amygdala ReHo and EPDS sum score at 12 months | Child sex, age at scan, maternal SES, maternal pre-pregnancy BMI, and ponderal index | 0.39 | < 0.01 |
| Left Amygdala ReHo and maternal pre-pregnancy BMI | Child sex, age at scan, maternal SES, ponderal index, and EPDS sum score at 6 months | -0.25 | < 0.05 |
| Left Amygdala ReHo and maternal pre-pregnancy BMI | Child sex, age at scan, maternal SES, ponderal index, EPDS sum score at 6 months, and SCL-90 sum score at 6 months | -0.25 | < 0.05 |

ReHo, regional homogeneity; EPDS, Edinburgh Postnatal Depression Scale; SES, socioeconomic status; BMI, body mass index; SCL-90, Symptom Checklist 90.

There was a significant positive association between left amygdala mean ReHo value and EPDS score at gestational week 24 when child sex and age at scan were controlled (*p* < 0.05, *β* = 0.25). This association did not persist when maternal SES, maternal pre-pregnancy BMI, and ponderal index nor when SCL-90 score at gestational week 24 were statistically controlled for. There were no associations with the right amygdala mean ReHo value and EPDS score at gestational week 24.

We found a positive association between mean left amygdala mean ReHo value and EPDS score at 3 months postnatal when child sex and age at scan were included as covariates (*p* < 0.001, *β* = 0.48), and when maternal SES, maternal pre-pregnancy BMI and ponderal index were controlled in addition (*p* < 0.001, *β* = 0.47). When SCL-90 score at 3 months postnatal was additionally set as covariate the association attenuated (*p* < 0.05, *β* = 0.34). Right amygdala mean ReHo value and EPDS score at 3 months postnatal had statistically significant association when child sex and age at scan were controlled (*p* < 0.05, *β* = 0.28), and the association remained significant when additionally controlling for maternal SES, maternal pre-pregnancy BMI and ponderal index (*p* < 0.05, *β* = 0.29), but did not remain significant when SCL-90 score at 3 months postnatal was controlled in addition.

Left amygdala mean ReHo value and EPDS score at 6 months postnatal had a positive association when sex and age were controlled (*p* < 0.01, *β* = 0.38). The same association was found when maternal SES, maternal pre-pregnancy BMI and ponderal index were controlled in addition (*p* < 0.01, *β* = 0.37). The association did not remain significant when SCL-90 score at 6 months postnatal was controlled in addition. There were no associations with right amygdala mean ReHo value and EPDS score at 6 months. Lastly, when EPDS score at 6 months postnatal, child sex, age at scan, maternal SES, and ponderal index were controlled, we found association between left amygdala ReHo and maternal pre-pregnancy BMI (*p* < 0.05, *β* = −0.25) and the association remained when SCL-90 score at 6 months postnatal was controlled in addition.

There was statistically significant association between left amygdala mean ReHo value and EPDS score at 12 months postnatal when child sex and age at scan were controlled (*p* < 0.001, *β* = 0.42), but did not persist when maternal SES, maternal pre-pregnancy BMI and ponderal index were controlled in addition (*p* < 0.01, *β* = 0.39). No associations between right amygdala mean ReHo values and EPDS score at 12 months postnatal were found.

## 4. Discussion

In this study, we investigated whether maternal perinatal depressive symptoms associate with child’s bilateral amygdala ReHo and seed-based functional connectivity. The voxel-wise analyses of ReHo implicated that the left amygdala had a strong positive association to EPDS scores obtained 3 months postnatally, and similar associations were detected for EPDS scores at 6 and 12 months postnatal albeit with smaller effect sizes. Association between left amygdala ReHo values and depressive symptoms did not persist when maternal anxiety symptoms were controlled in the whole brain analysis but were the strongest predictor in the ROI analyses (Supplemental A). Seed-based connectivity analysis did not reveal any significant associations in the connectivity of the left amygdala and the rest of the brain. We were thus unable to replicate prior fundings (13).

Our study showed that perinatal maternal depressive symptoms were associated with heightened ReHo, i.e. heightened local functional activity, in the offspring’s left amygdala. These findings are intriguing given that the association was relatively strong and particularly localized in the left amygdala in the whole brain statistical models. Research of the mechanisms underlying depression have shown that left amygdala hyperactivity increases the risk for depression (27–29). Considering the above-mentioned and our findings, which suggest that the offspring of mothers experiencing perinatal depressive symptoms have heightened left amygdala functional synchrony that implicates either synchronized lower or heightened brain activity. It raises the possibility that amygdala regional connectivity can be a neural marker of intergenerational transmission of risk for depression and other affective disorders. However, this remains to be addressed in future studies.

The effect of postnatal maternal depressive symptoms on child’s developing brain mediates via parenting and mother–child interaction (30,31). Infancy and toddlerhood are crucial periods for the development of child’s self- and emotion-regulation, cognitive, language and motor skills (32–34). A key feature of depression is dysfunction of self- and emotion-regulation (35). Therefore, maternal depressive symptoms may affect the self-regulation and sensitivity of the mother. Sensitive interaction refers to mother’s ability to interpret child’s physical needs and emotional cues and to response appropriately (36). Preschool-aged children of mothers experiencing postnatal depression had more internalizing and externalizing behavioural problems, i.e. altered self- and emotion-regulation (37,38).

Furthermore, postnatal depression has been found to be negatively associated with white matter integrity in vast areas in 5-year-old girls (39).

### 4.1 Limitations

Although our sample size is relatively small, our results are compatible with the few previous studies investigating similar effects using fMRI (9,13). Nevertheless, studies with larger cohorts are needed in the future. The study population’s cultural background was narrow and therefore it is needed to further study this association in other populations. As the children were scanned at one time point, further studies are needed to determine whether amygdala functional changes persist or evolve. Regarding the limitations of fMRI acquisition, the use of a passive viewing paradigm to reduce head motion (40,41) and the presence of the parent in the imaging room may have affected the measured fMRI signal. The EPDS and SCL-90 scores are self-reported measures and majority of the participating mothers did not have clinically meaningful or severe symptoms of perinatal depression.

### 4.2 Conclusions

In conclusion, we found that maternal perinatal depressive symptoms were associated with heightened local functional connectivity of the left amygdala in 5-year-olds. We did not find associations in the patterns of the left amygdala connectivity to the rest of the brain. Our results provide preliminary indication that maternal mental health during pregnancy and especially soon after birth may influence offspring’s neurodevelopment related to emotional regulation.

## Acknowledgements

We would like to warmly thank all FinnBrain families that participated to the study. We would also like to thank the research team, Susanne Sinisalo for performing the MRI scans with the investigators, Jani Saunavaara for implementing the MRI sequences, Riitta Parkkola for reviewing the MR images for incidental findings, and Tuire Lähdesmäki for being the consulting pediatric neurologist.

## 6.2 Author contributions

- Elena Vartiainen, performed seed-based connectivity pre-processing, performed the statistical models and lead the writing of the manuscript.
- Anni Copeland, collected the MRI data, fMRI data quality control and pre-processing, performed the statistical models, wrote the initial draft of manuscript and supervised Elena Vartiainen.
- Elmo P. Pulli collected the MRI data and participated in the initial draft of manuscript, supervised Elena Vartiainen.
- Venla Kumpulainen, collected the MRI data.
- Eero Silver, collected the MRI data.
- Olli Rajasilta, supported the data preprocessing and analyses.
- Ashmeet Jolly, participated in the initial draft of the manuscript.
- Silja Luotonen, participated in the initial draft of the manuscript.
- Hilyatushalihah K. Audah, participated in the initial draft of the manuscript.
- Niloofar Hashempour, participated in the initial draft of the manuscript.
- Wajiha Bano, participated in the initial draft of the manuscript.
- Ilkka Suuronen, participated in the initial draft of the manuscript.
- Ekaterina Saukko, collected the MRI data.
- Suvi Häkkinen, designed the data processing pipelines.
- Hasse Karlsson, conceptualized the study and build the infrastructure of the FinnBrain Birth Cohort
- Linnea Karlsson, conceptualized the study and build the infrastructure of the FinnBrain Birth Cohort.
- Jetro J. Tuulari conceptualized the study, planned, and funded the measurements, designed the data processing pipelines, supervised Elena Vartiainen and Anni Copeland.
- All co-authors participated in editing the manuscript and accepted it in its final form.

## Supplemental table and figure legends

**Supplemental Table A.** Regression results for the ReHo of the Left Amygdala.

**Supplemental Table B.** Regression results for the ReHo of the Right Amygdala.

**Supplemental Figure 1.** ReHo values of the left amygdala associate positively with EPDS scores at 3 months postnatal; controlling for child sex and age at scan. The results have been thresholded at p < 0.001, FDR multiple comparisons corrected at the cluster level. The color bars depict t-values.

