## Supplements for "Pre- and postnatal maternal depressive symptoms associate with local connectivity of the left amygdala in 5-year-olds"

**Supplemental Table A.** Regression results using the ReHo of the Left Amygdala as the criterion.

| Left amygdala and EPDS sum score at gestational week 24 | | | | | | | | |
| --- | --- | --- | --- | --- | --- | --- | --- | --- |
| Predictor | *b* | *b*  95% CI  [LL, UL] | *beta* | *beta*  95% CI  [LL, UL] | *sr^2^* | *sr^2^*  95% CI  [LL, UL] | *r* | Fit |
| (Intercept) | 1.07 | [-0.29, 2.43] |  |  |  |  |  |  |
| EPDS sum score at GW 24 | 0.00 | [-0.01, 0.01] | 0.10 | [-0.26, 0.45] | 0.00 | [-.02, .03] | 0.27* |  |
| Child sex | 0.03 | [-0.03, 0.10] | 0.14 | [-0.12, 0.39] | 0.02 | [-.04, .07] | 0.12 |  |
| Age at scan | -0.00 | [-0.00, 0.00] | -0.07 | [-0.31, 0.17] | 0.01 | [-.03, .04] | -0.07 |  |
| SES | -0.02 | [-0.06, 0.01] | -0.15 | [-0.39, 0.09] | 0.02 | [-.04, .08] | -0.11 |  |
| Pre-pregnancy BMI | -0.01 | [-0.02, 0.00] | -0.25 | [-0.50, 0.01] | 0.05 | [-.05, .15] | -0.18 |  |
| Ponderal index | 0.02 | [-0.01, 0.04] | 0.15 | [-0.11, 0.40] | 0.02 | [-.04, .08] | 0.08 |  |
| SCL sum score at GW 24 | 0.01 | [-0.01, 0.02] | 0.17 | [-0.18, 0.51] | 0.01 | [-.04, .06] | 0.23 |  |
|  |  |  |  |  |  |  |  | *R^2^* = .158  95% CI[.00,.24] |
| Left amygdala and EPDS sum score at 3 months | | | | | | | | |
| Predictor | *b* | *b*  95% CI  [LL, UL] | *beta* | *beta*  95% CI  [LL, UL] | *sr^2^* | *sr^2^*  95% CI  [LL, UL] | *r* | Fit |
| (Intercept) | 0.54 | [-0.70, 1.79] |  |  |  |  |  |  |
| EPDS sum score at 3 months | 0.01* | [0.00, 0.02] | 0.34 | [0.04, 0.63] | 0.06 | [-.03, .15] | 0.48** | <0.05 |
| Child sex | 0.04 | [-0.02, 0.09] | 0.15 | [-0.06, 0.37] | 0.02 | [-.04, .08] | 0.12 |  |
| Age at scan | 0.00 | [-0.00, 0.00] | 0.01 | [-0.21, 0.24] | 0.00 | [-.01, .01] | -0.07 |  |
| SES | -0.02 | [-0.05, 0.01] | -0.13 | [-0.35, 0.09] | 0.02 | [-.03, .06] | -0.11 |  |
| Pre-pregnancy BMI | -0.01 | [-0.01, 0.00] | -0.19 | [-0.41, 0.04] | 0.03 | [-.04, .10] | -0.18 |  |
| Ponderal index | 0.01 | [-0.01, 0.04] | 0.12 | [-0.10, 0.35] | 0.01 | [-.03, .06] | 0.08 |  |
| SCL sum score at 3 months | 0.01 | [-0.00, 0.02] | 0.19 | [-0.11, 0.49] | 0.02 | [-.03, .07] | 0.46** |  |
|  |  |  |  |  |  |  |  | *R^2^* = .325**  95% CI[.07,.42] |
| Left amygdala and EPDS sum score at 6 months | | | | | | | | |
| Predictor | *b* | *b*  95% CI  [LL, UL] | *beta* | *beta*  95% CI  [LL, UL] | *sr^2^* | *sr^2^*  95% CI  [LL, UL] | *r* | Fit |
| (Intercept) | 0.97 | [-0.33, 2.27] |  |  |  |  |  |  |
| EPDS sum score at 6 months | 0.01 | [-0.00, 0.02] | 0.30 | [-0.14, 0.73] | 0.02 | [-.04, .09] | 0.39** |  |
| Child sex | 0.03 | [-0.03, 0.09] | 0.12 | [-0.11, 0.35] | 0.01 | [-.03, .06] | 0.12 |  |
| Age at scan | -0.00 | [-0.00, 0.00] | -0.06 | [-0.29, 0.17] | 0.00 | [-.02, .03] | -0.07 |  |
| SES | -0.02 | [-0.06, 0.01] | -0.14 | [-0.37, 0.09] | 0.02 | [-.04, .07] | -0.11 |  |
| Pre-pregnancy BMI | -0.01* | [-0.02, -0.00] | -0.25 | [-0.49, -0.01] | 0.05 | [-.04, .15] | -0.18 |  |
| Ponderal index | 0.02 | [-0.01, 0.04] | 0.15 | [-0.09, 0.39] | 0.02 | [-.04, .08] | 0.08 |  |
| SCL sum score at 6 months | 0.00 | [-0.01, 0.01] | 0.09 | [-0.35, 0.52] | 0.00 | [-.02, .02] | 0.36** |  |
|  |  |  |  |  |  |  |  | *R^2^* = .237*  95% CI[.01,.33] |
| Left amygdala and EPDS sum score at 12 months | | | | | | | | |
| Predictor | *b* | *b*  95% CI  [LL, UL] | *beta* | *beta*  95% CI  [LL, UL] | *sr^2^* | *sr^2^*  95% CI  [LL, UL] | *r* | Fit |
| (Intercept) | 0.80 | [-0.49, 2.09] |  |  |  |  |  |  |
| EPDS sum score at 12 months | 0.01** | [0.00, 0.02] | 0.39 | [0.16, 0.61] | 0.14 | [-.00, .29] | 0.42** |  |
| Child sex | 0.04 | [-0.02, 0.09] | 0.15 | [-0.08, 0.38] | 0.02 | [-.04, .08] | 0.12 |  |
| Age at scan | -0.00 | [-0.00, 0.00] | -0.03 | [-0.26, 0.20] | 0.00 | [-.01, .01] | -0.07 |  |
| SES | -0.02 | [-0.05, 0.01] | -0.13 | [-0.36, 0.10] | 0.02 | [-.04, .07] | -0.11 |  |
| Pre-pregnancy BMI | -0.01 | [-0.01, 0.00] | -0.20 | [-0.44, 0.04] | 0.03 | [-.04, .11] | -0.18 |  |
| Ponderal index | 0.01 | [-0.01, 0.04] | 0.13 | [-0.11, 0.37] | 0.01 | [-.03, .06] | 0.08 |  |
|  |  |  |  |  |  |  |  | *R^2^* = .242**  95% CI[.02,.35] |

*Note.* A significant *b*-weight indicates the beta-weight and semi-partial correlation are also significant. *b* represents unstandardized regression weights. *beta* indicates the standardized regression weights. *sr^2^* represents the semi-partial correlation squared. *r* represents the zero-order correlation. *LL* and *UL* indicate the lower and upper limits of a confidence interval, respectively.
* indicates *p*<0.05. ** indicates *p*<0.01.

EPDS, Edinburgh Postnatal Depression Scale; GW, gestational week; SES, socioeconomic status; pre-pregnancy BMI, maternal pre-pregnancy body mass index; SCL, Symptom Checklist.

**Supplemental Table B.** Regression results using the ReHo of the Right Amygdala as the criterion.

| Right amygdala and EPDS sum score at gestational week 24 | | | | | | | | |
| --- | --- | --- | --- | --- | --- | --- | --- | --- |
| Predictor | *b* | *b*  95% CI  [LL, UL] | *beta* | *beta*  95% CI  [LL, UL] | *sr^2^* | *sr^2^*  95% CI  [LL, UL] | *r* | Fit |
| (Intercept) | 1.54* | [0.06, 3.02] |  |  |  |  |  |  |
| EPDS sum score GW 24 | 0.00 | [-0.01, 0.02] | 0.16 | [-0.22, 0.53] | .01 | [-.04, .06] | .15 |  |
| Child sex | -0.01 | [-0.08, 0.05] | -0.06 | [-0.33, 0.21] | .00 | [-.02, .03] | -.05 |  |
| Age at scan | -0.00 | [-0.00, 0.00] | -0.15 | [-0.41, 0.10] | .02 | [-.05, .09] | -.17 |  |
| SES | -0.01 | [-0.05, 0.03] | -0.06 | [-0.32, 0.20] | .00 | [-.02, .03] | -.07 |  |
| Pre-pregnancy BMI | 0.00 | [-0.01, 0.01] | 0.06 | [-0.20, 0.33] | .00 | [-.02, .03] | .06 |  |
| Ponderal index | 0.00 | [-0.03, 0.03] | 0.03 | [-0.24, 0.30] | .00 | [-.01, .02] | .04 |  |
| SCL sum score at GW 24 | 0.00 | [-0.01, 0.01] | 0.01 | [-0.36, 0.37] | .00 | [-.00, .00] | .13 |  |
|  |  |  |  |  |  |  |  | *R^2^* = .064  95% CI[.00,.10] |
| Right amygdala and EPDS sum score at 3 months | | | | | | | | |
| Predictor | *b* | *b*  95% CI  [LL, UL] | *beta* | *beta*  95% CI  [LL, UL] | *sr^2^* | *sr^2^*  95% CI  [LL, UL] | *r* | Fit |
| (Intercept) | 1.27 | [-0.20, 2.74] |  |  |  |  |  |  |
| EPDS sum score at 3 months | 0.01 | [-0.00, 0.02] | 0.32 | [-0.02, 0.66] | .05 | [-.05, .15] | .31* |  |
| Child sex | -0.01 | [-0.07, 0.06] | -0.03 | [-0.28, 0.22] | .00 | [-.01, .01] | -.05 |  |
| Age at scan | -0.00 | [-0.00, 0.00] | -0.11 | [-0.36, 0.15] | .01 | [-.03, .06] | -.17 |  |
| SES | -0.01 | [-0.05, 0.03] | -0.08 | [-0.33, 0.17] | .01 | [-.03, .04] | -.07 |  |
| Pre-pregnancy BMI | 0.00 | [-0.01, 0.01] | 0.07 | [-0.19, 0.33] | .00 | [-.03, .03] | .06 |  |
| Ponderal index | 0.00 | [-0.03, 0.03] | 0.02 | [-0.24, 0.28] | .00 | [-.01, .01] | .04 |  |
| SCL sum score at 3 months | -0.00 | [-0.02, 0.01] | -0.04 | [-0.39, 0.30] | .00 | [-.01, .01] | .20 |  |
|  |  |  |  |  |  |  |  | *R^2^* = .120  95% CI[.00,.19] |
| Left amygdala and EPDS sum score at 6 months | | | | | | | | |
| Predictor | *b* | *b*  95% CI  [LL, UL] | *beta* | *beta*  95% CI  [LL, UL] | *sr^2^* | *sr^2^*  95% CI  [LL, UL] | *r* | Fit |
| (Intercept) | 1.60* | [0.14, 3.06] |  |  |  |  |  |  |
| EPDS sum score at 6 months | 0.01 | [-0.00, 0.03] | 0.43 | [-0.04, 0.90] | .05 | [-.05, .15] | .20 |  |
| Child sex | -0.01 | [-0.07, 0.06] | -0.03 | [-0.29, 0.22] | .00 | [-.01, .02] | -.05 |  |
| Age at scan | -0.00 | [-0.00, 0.00] | -0.16 | [-0.41, 0.09] | .02 | [-.05, .09] | -.17 |  |
| SES | -0.01 | [-0.05, 0.03] | -0.06 | [-0.31, 0.19] | .00 | [-.02, .03] | -.07 |  |
| Pre-pregnancy BMI | 0.00 | [-0.01, 0.01] | 0.03 | [-0.23, 0.29] | .00 | [-.01, .01] | .06 |  |
| Ponderal index | 0.00 | [-0.03, 0.03] | 0.02 | [-0.24, 0.29] | .00 | [-.01, .01] | .04 |  |
| SCL sum score at 6 months | -0.01 | [-0.02, 0.00] | -0.29 | [-0.77, 0.18] | .02 | [-.05, .09] | .09 |  |
|  |  |  |  |  |  |  |  | *R^2^* = .097  95% CI[.00,.16] |
| Right amygdala and EPDS sum score at 12 months | | | | | | | | |
| Predictor | *b* | *b*  95% CI  [LL, UL] | *beta* | *beta*  95% CI  [LL, UL] | *sr^2^* | *sr^2^*  95% CI  [LL, UL] | *r* | Fit |
| (Intercept) | 1.50* | [0.01, 2.99] |  |  |  |  |  |  |
| EPDS sum score at 12 months | 0.00 | [-0.00, 0.01] | 0.13 | [-0.13, 0.38] | .02 | [-.04, .07] | .15 |  |
| Child sex | -0.01 | [-0.07, 0.06] | -0.03 | [-0.28, 0.23] | .00 | [-.01, .01] | -.05 |  |
| Age at scan | -0.00 | [-0.00, 0.00] | -0.14 | [-0.40, 0.11] | .02 | [-.04, .08] | -.17 |  |
| SES | -0.01 | [-0.05, 0.03] | -0.06 | [-0.32, 0.19] | .00 | [-.02, .03] | -.07 |  |
| Pre-pregnancy BMI | 0.00 | [-0.01, 0.01] | 0.07 | [-0.20, 0.33] | .00 | [-.02, .03] | .06 |  |
| Ponderal index | 0.00 | [-0.03, 0.03] | 0.03 | [-0.24, 0.29] | .00 | [-.01, .01] | .04 |  |
|  |  |  |  |  |  |  |  | *R^2^* = .056  95% CI[.00,.10] |

*Note.* A significant *b*-weight indicates the beta-weight and semi-partial correlation are also significant. *b* represents unstandardized regression weights. *beta* indicates the standardized regression weights. *sr^2^* represents the semi-partial correlation squared. *r* represents the zero-order correlation. *LL* and *UL* indicate the lower and upper limits of a confidence interval, respectively.
* indicates *p*<0.05. ** indicates *p*<0.01.

EPDS, Edinburgh Postnatal Depression Scale; GW, gestational week; SES, socioeconomic status; pre-pregnancy BMI, maternal pre-pregnancy body mass index; SCL, Symptom Checklist.

**Supplemental Figure 1.** ReHo values of the left amygdala associate positively with EPDS scores at 3 months postnatal; controlling for child sex and age at scan. The results have been thresholded at p < 0.001, FDR multiple comparisons corrected at the cluster level. The color bars depict t-values.

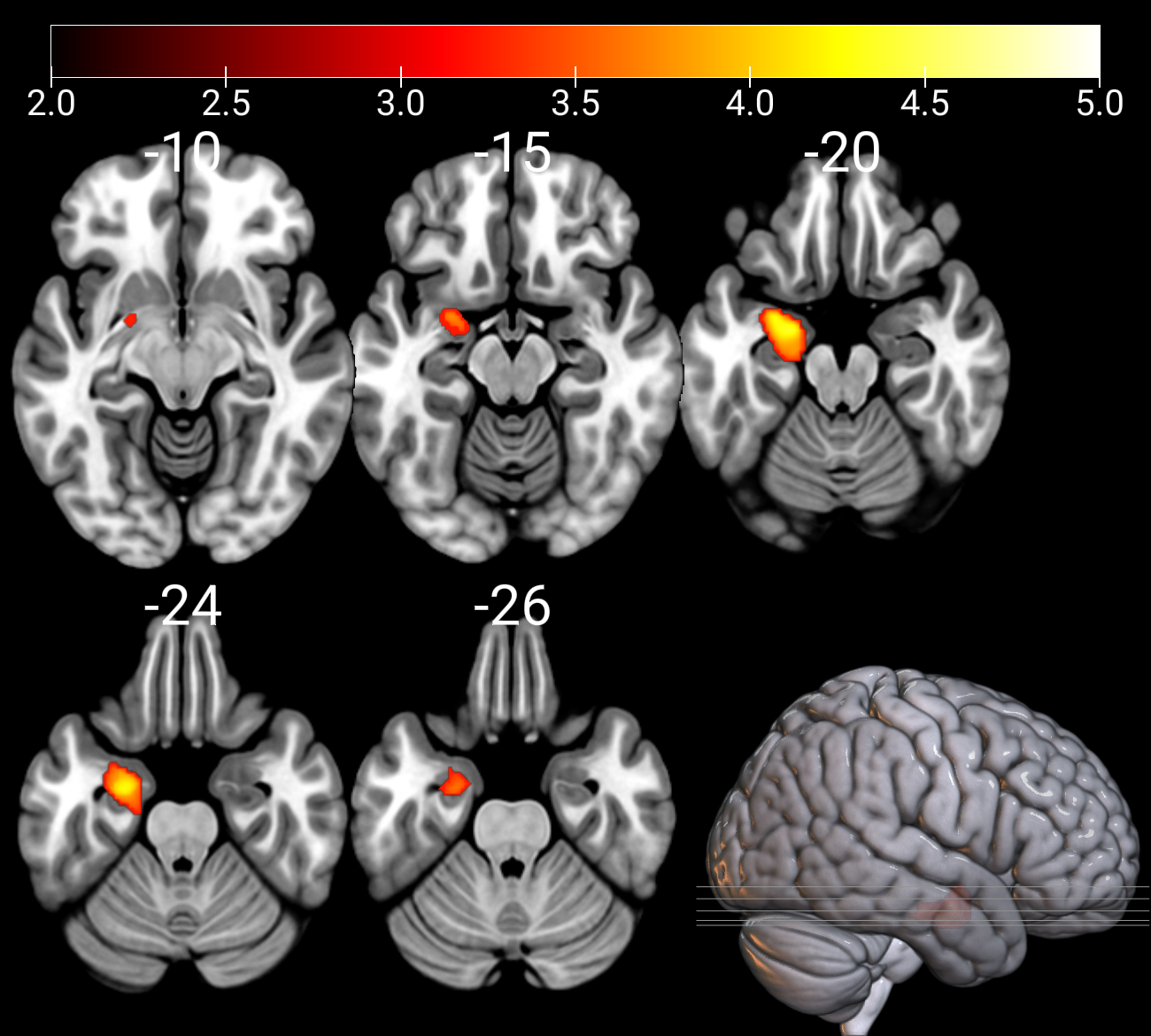
